# Assessing the prevalence of mycoplasma contamination in cell culture via a survey of NCBI’s RNA-seq archive

**DOI:** 10.1101/007054

**Authors:** Anthony O. Olarerin-George, John B. Hogenesch

## Abstract

Mycoplasmas are notorious contaminants of cell culture and can have profound effects on host cell biology by depriving cells of nutrients and inducing global changes in gene expression. Because they are small, they can escape filtration in culture media. Because they lack cell walls, they are resistant to commonly used antibiotics. Over the last two decades, sentinel testing has revealed wide-ranging contamination rates in mammalian culture. To obtain an unbiased assessment from hundreds of labs, we analyzed sequence data from 9395 rodent and primate samples from 884 series (or projects) in the NCBI Sequence Read Archive. We found 11% of these series were contaminated (defined as ≥ 100 reads/million mapping to mycoplasma in one or more samples). Ninety percent of mycoplasma-mapped reads aligned to ribosomal RNA. Interestingly, series using poly(A)-selection, which should bias against mycoplasma detection, had comparable contamination rates as non-poly(A)-selected series. We also examined the relationship between mycoplasma contamination and host gene expression in a single cell RNA-seq dataset and found 61 host genes (*P* < 0.001) were significantly associated with mycoplasma-mapped read counts. Lastly, to estimate the potential economic cost of this widespread contamination, we queried NIH RePORTER to find grants with the terms “cell culture” or “cell lines”. Funding for these totaled over $3 billion, suggesting hundreds of millions of dollars in research are potentially affected. In all, this study suggests mycoplasma contamination is still prevalent today and poses substantial risk to research quality, with considerable financial consequences.

## INTRODUCTION

Mycoplasmas are small parasitic bacteria of the class mollicutes. There are over 180 species infecting a wide range of hosts. Mycoplasmas are common to the human respiratory and urogenital tracts (Taylor-Robinson 1996). Some species are pathogenic. For example, *M. pneumoniae* causes atypical pneumonia (Chanock et al. 1962). Also, *M. genitalium* infection is linked to pelvic inflammatory disease (Tully et al. 1981).

In addition to their impact on human health, mycoplasmas are widespread contaminants of cell culture. In 1956, researchers from Johns Hopkins reported mycoplasma contamination of HeLa cells used in their lab (Robinson and Wichelhausen 1956). By the early 1990s, the US Food and Drug Administration had tested over 20,000 cell cultures and found 15% contaminated with mycoplasma (Rottem and Barile 1993). A study in 1991 from Argentina found 70% of the 200 samples tested were contaminated (Coronato and Coto 1991). More recently, a 2002 report by the Deutsche Sammlung von Mikroorganismen und Zellkulturen (DSMZ) in Germany found 28% of the 440 cell lines tested (mostly leukemia-lymphoma) were contaminated (Drexler and Uphoff 2002). Hence, mycoplasma contamination in cell culture is a long-standing and persistent problem.

Preventing mycoplasma contamination is difficult. For one, mycoplasma cells are small (0.3-0.8 uM in diameter) and pleomorphic (Razin 2006), allowing them to pass through standard filtration membranes. Secondly, mycoplasmas lack cell walls. This makes them impervious to cell culture antibiotics that inhibit cell wall synthesis, such as penicillin. Effective antibiotics do exist, however, their continuous use in cell culture is not recommended due to possible cytotoxicity. Third, mycoplasmas are able to reach high concentrations in the media of infected cells without noticeable turbidity (Young et al. 2010). This makes detection via visual inspection difficult. Lastly, while the primary source of mycoplasma contamination is likely other cell cultures, mycoplasmas from lab personnel are a potential source as well (Drexler and Uphoff 2002).

Mycoplasmas have one of the smallest prokaryotic genomes (∼0.6 Mb). This comes at a price, as mycoplasmas lack key genes essential for the synthesis of macromolecule precursors and energy metabolism (Razin et al. 1998). As a result, mycoplasmas alter and depend on host cell biology for survival (Rottem and Naot 1998; Rottem 2003). For example, *M. orale* can compete for arginine in culture media, impacting host cell growth (Gill and Pan 1970). Also, *M. hyorhinis* endonucleases can degrade host cell DNA, providing DNA precursors for the parasite (Paddenberg et al. 1998). Further, mycoplasma infection can dysregulate hundreds of host genes (Hopfe et al. 2013). It is therefore imperative that cultured cells are free of mycoplasma contamination to ensure the interpretability, reproducibility and reliability of results obtained from their use.

In this study we sought to evaluate the prevalence of mycoplasma contamination in cell culture today. High throughput RNA-sequencing data is growing at an exponential rate and is providing an unprecedented view of the constituency of RNA molecules in a sample. We posited that mycoplasma sequences in RNA-Seq data from primate and rodent specimens would be indicative of contamination. Further, because these samples come from hundreds of geographically distinct labs, it offers a more comprehensive view of contamination. Hence, we surveyed RNA-Seq data from archives at NCBI for mycoplasma sequences. We also evaluated the relationship between mycoplasma contamination and host gene expression in a Burkitt’s lymphoma cell line.

## RESULTS

### Acquisition and characterization of RNA-Seq data

We assessed the feasibility of working with the large volume of sequencing data in NCBI’s Sequence Read Archive (SRA). At the time of this study, there were 884 series (or projects) of type “expression profiling by high throughput sequencing” or “non-coding RNA profiling by high throughput sequencing” from primates and rodents (**Supplemental data 1**). These were comprised of 9395 samples with downloadable sequence files. To make the analysis manageable, we downloaded just the first million reads of each of the samples using NCBI’s SRA toolkit. For samples with paired-end reads, we assessed only the first read. The resulting dataset contained 6.4 billion reads and filled 2TB of disk space.

### Mapping of reads to mycoplasma

We mapped the reads to mycoplasma in two steps. First, we aligned the reads to four complete mycoplasma genomes (*M. hominis* ATCC 23114, *M. hyorhinis* MCLD, *M. fermentans* M64 and *A. laidlawii* PG-8A) with Bowtie (Langmead et al. 2009). Less than 2% (112,547,232) of the reads mapped to one or more of the mycoplasma species. This corresponded to 1,280,951 unique sequence reads (**Supplemental data 2**). Next, we eliminated non-specific bowtie aligned reads. These were reads that mapped to sequences from other sources as well as or better than they did mycoplasma. For example, reads that mapped to the putative host species or to other bacteria were eliminated. To accomplish this, we aligned the unique bowtie-mapped reads to NCBI’s nucleotide collection (nt) database with standalone BLAST+ (Altschul et al. 1990). The nt database is a comprehensive collection of ∼20 million DNA sequences from species spanning the phylogeny of life and viruses. These sequences include entries from Refseq, GenBank, EMBL and DDBJ. Of the 1,280,951 unique bowtie aligned reads, 474,219 (37%) aligned best to mycoplasma as assessed by BLAST E-values (**Supplemental Table 1**). Ninety percent of all BLAST-confirmed reads aligned to ribosomal sequences (Figure 1).

**Figure 1.**
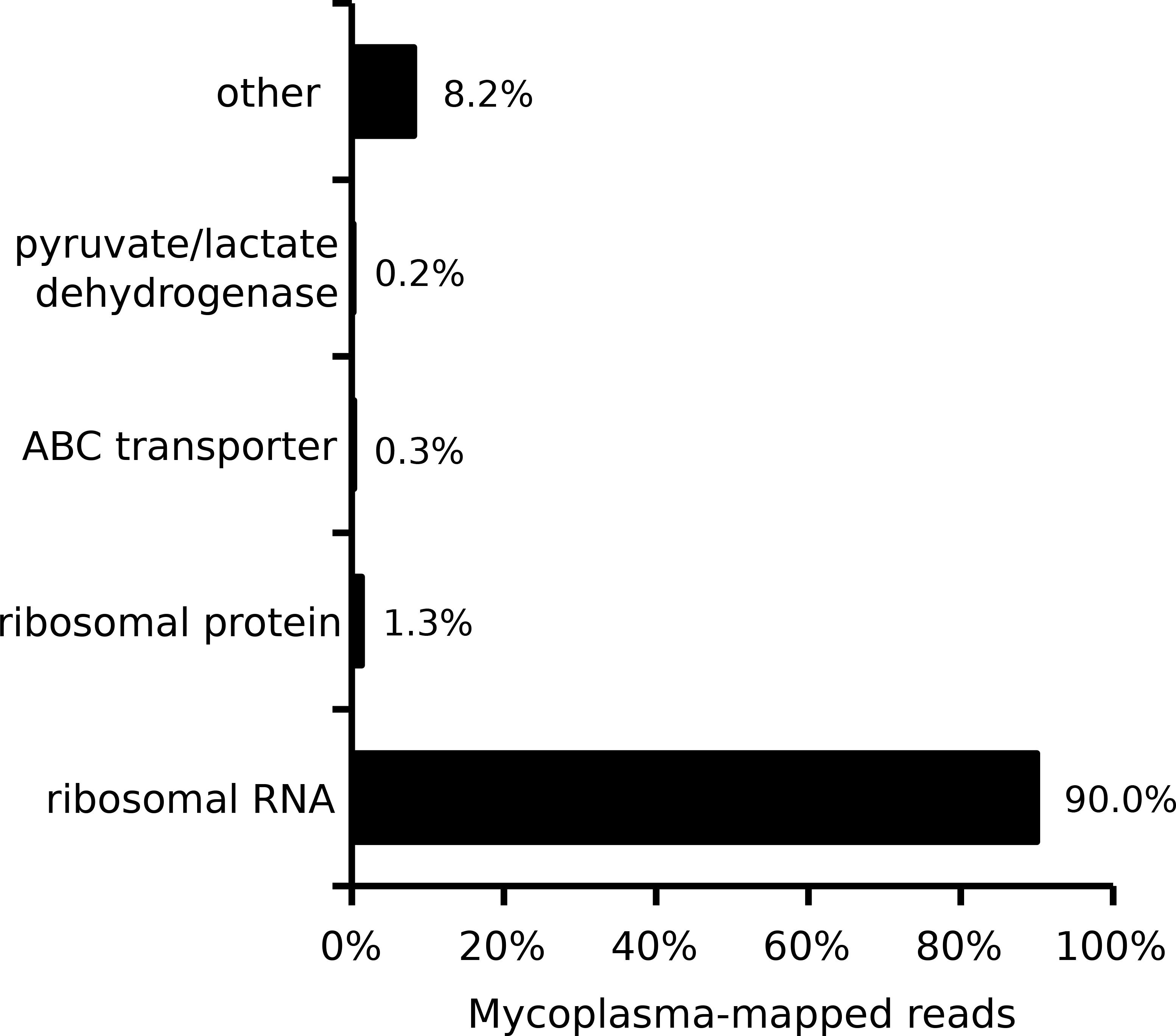
Gene breakdown of mycoplasma-mapped reads. RNA-seq reads were aligned to four mycoplasma genomes using bowtie. Non-specific reads were filtered with BLAST. Of the resulting 472,219 mycoplasma-mapped reads, 90% mapped to ribosomal RNA.

### Mycoplasma contamination in cultured vs non-cultured cells

Mycoplasma contamination is predominantly found in cultured samples (e.g. cell lines) and not in non-cultured samples (e.g. tissues) (Miller et al. 2003). Hence, we posited that non-cultured samples in our study would be underrepresented for mycoplasma contamination and provide a baseline for the expected number of mycoplasma-mapped reads in a given sample (i.e. serve as a negative control). We mined the text of the sample descriptions from NCBI’s Gene Expression Omnibus (GEO) for keywords indicating if the samples were cultured or not (see Methods). We estimated that 57% of the samples (n = 5328) were cultured and 43% (n = 4067) were not (Supplemental Table 2). In series with cultured samples, 11% (52/484) had at least one sample with 100 or more reads per million (RPM) that mapped to mycoplasma (see full distribution in Figure 2A). In contrast, none of the 344 series with non-cultured samples met this criterion (Figure 2A). Further, more samples had reads mapping to *M. hyorhinis,* than the other tested mycoplasma species (Figure 2B, 2C). Fewer samples had reads mapping to *A. laidlawii*. Surprisingly, for 39 samples (all cultured), more than 10^4^ RPM (i.e >1% of reads) mapped to one or more of the mycoplasma species (Figure 2A, **Supplemental Table 2**).

**Figure 2.**
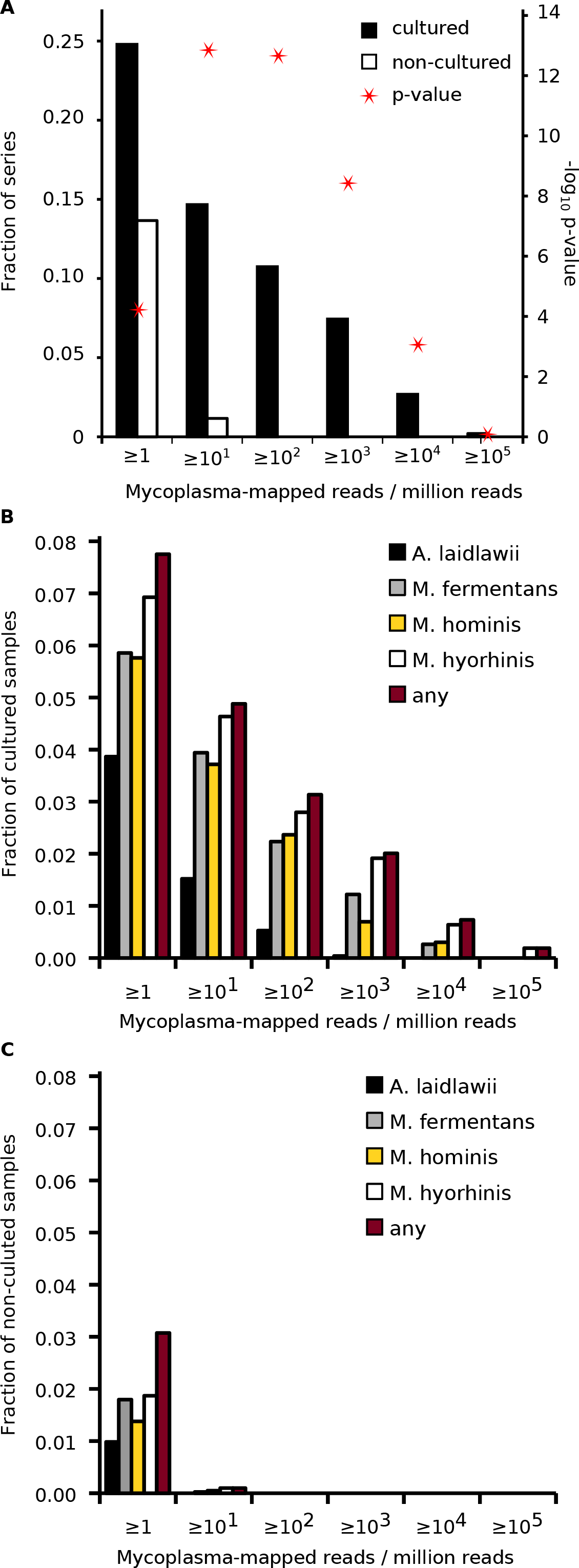
Mycoplasma contamination in cultured vs non-cultured samples and series. A) Fraction of contaminated series (containing cultured samples or not) for various cutoffs of mycoplasma-mapped reads per million (column graphs; primary y-axis). Red stars indicate the p-values of the respective comparisons (secondary y-axis; Fisher’s exact test). Fraction of contaminated B) cultured or C) non-cultured samples for various cutoffs of mycoplasma-mapped reads per million, broken down by the indicated mycoplasma species.

### Mycoplasma contamination in poly(A)- vs non-poly(A)-selected samples

Polyadenylation marks transcripts for degradation in bacteria (Dreyfus and Régnier 2002, 9). Hence, we hypothesized that poly(A)-selected samples would be underrepresented for mycoplasma contamination. We mined the text of the sample descriptions from NCBI’s GEO for keywords indicating if the samples were poly(A)-selected or not (see Methods). We estimated that 26% (n = 2431) of the samples were poly(A) selected and 74% (n = 6964) were not (Supplemental Table 2). The latter category was broad and included samples that were depleted of ribosomal RNA, size-selected for small RNAs, immunoprecipitated via RNA binding proteins and others. Unexpectedly, there was no significant difference in the fraction of series with mycoplasma-mapped reads in the poly(A) and non-poly(A) series (Figure 3A). In general, more samples had reads mapping to *M. hyorhinis* than the other mycoplasma species. Fewer samples had reads mapping to *A. laidlawii* (Figure 3B, 3C).

**Figure 3.**
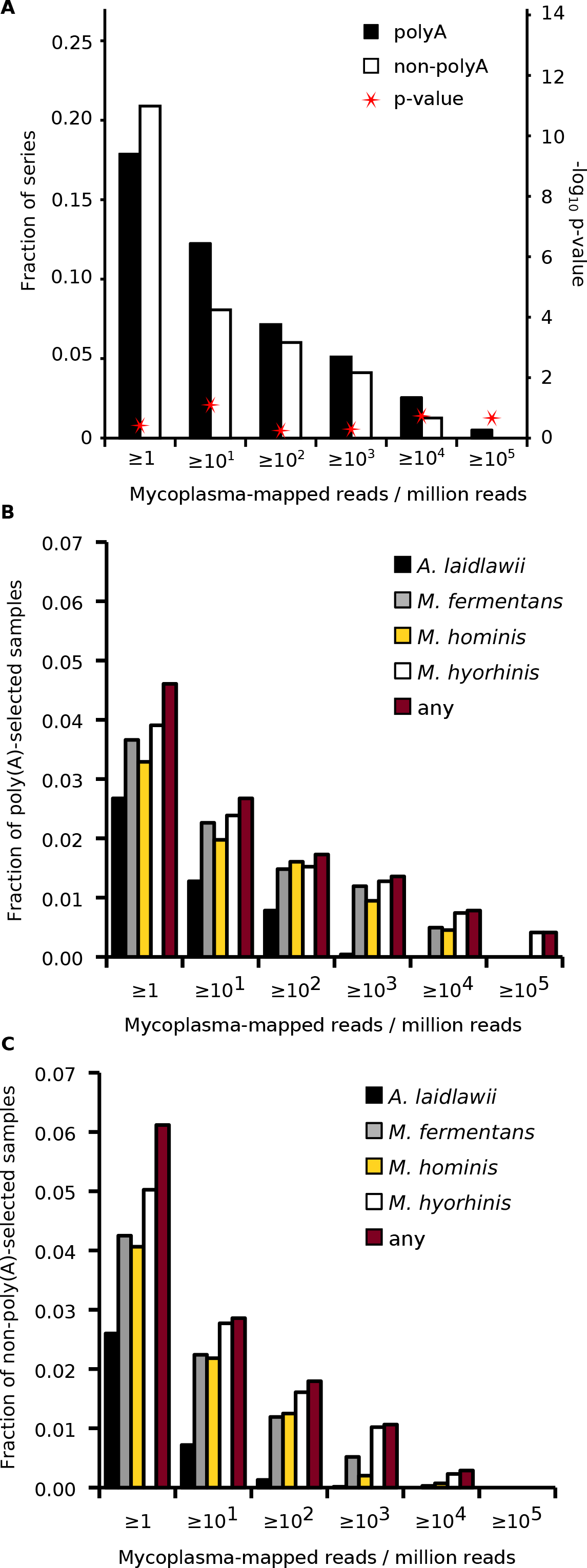
Mycoplasma contamination in poly(A)- vs non-poly(A)-selected samples and series. A) Fraction of contaminated series (containing poly(A)-selected samples or not) for various cutoffs of mycoplasma-mapped reads per million (column graphs; primary y-axis). Red stars indicate the p-values of the respective comparisons (secondary y-axis; Fisher’s exact test). Fraction of contaminated B) poly(A)- or C) non-poly(A)-selected samples for various cutoffs of mycoplasma-mapped reads per million, broken down by the indicated mycoplasma species.

### Effect of mycoplasma contamination on host gene expression

Next, we wanted to determine the effect of mycoplasma contamination on host gene expression. To do this, we searched for series that contained comparable contaminated and non-contaminated samples. This was rare as mycoplasma contamination in one sample typically guaranteed contamination of all samples of the same cell line/type in the series. Hence, we instead focused on contaminated series with large numbers of replicate samples and varied amounts of mycoplasma-mapped reads. Our goal was to identify host genes whose expression levels were statistically associated with the number of mycoplasma-mapped reads. We found one such series (GSE49321) containing seven contaminated samples from the Burkitt’s lymphoma cell line DG-75. Interestingly, each sample represented expression data from a single cell. The number of mycoplasma-mapped reads ranged from ∼10,000 to 36,000 RPM (Figure 4A). We downloaded the full RNA-seq datasets for these samples, aligned them to the human reference genome using STAR (Dobin et al. 2013), and obtained read counts for genes using HTSeq (Anders et al. 2014). We then used DESeq2 (Love et al. 2014) to generate generalized linear models of host gene expression as a function of mycoplasma-mapped reads. Sixty-one genes were statistically associated with mycoplasma-mapped read counts (Wald test, Benjamini-Hochberg adjusted P < 0.001; **Supplemental Table 3**). We found no significant enrichment in gene annotations using DAVID (Huang et al. 2009). The top 16 genes are shown in Figure 4A. Finally, we evaluated the likelihood of obtaining significant genes by chance in our analysis. We randomly permuted the gene expression counts of the samples and repeated the analysis 1000 times. The number of significant genes was consistently less than that of the observed set for various p-value cutoffs examined (P < 0.002, Figure 4B).

**Figure 4.**
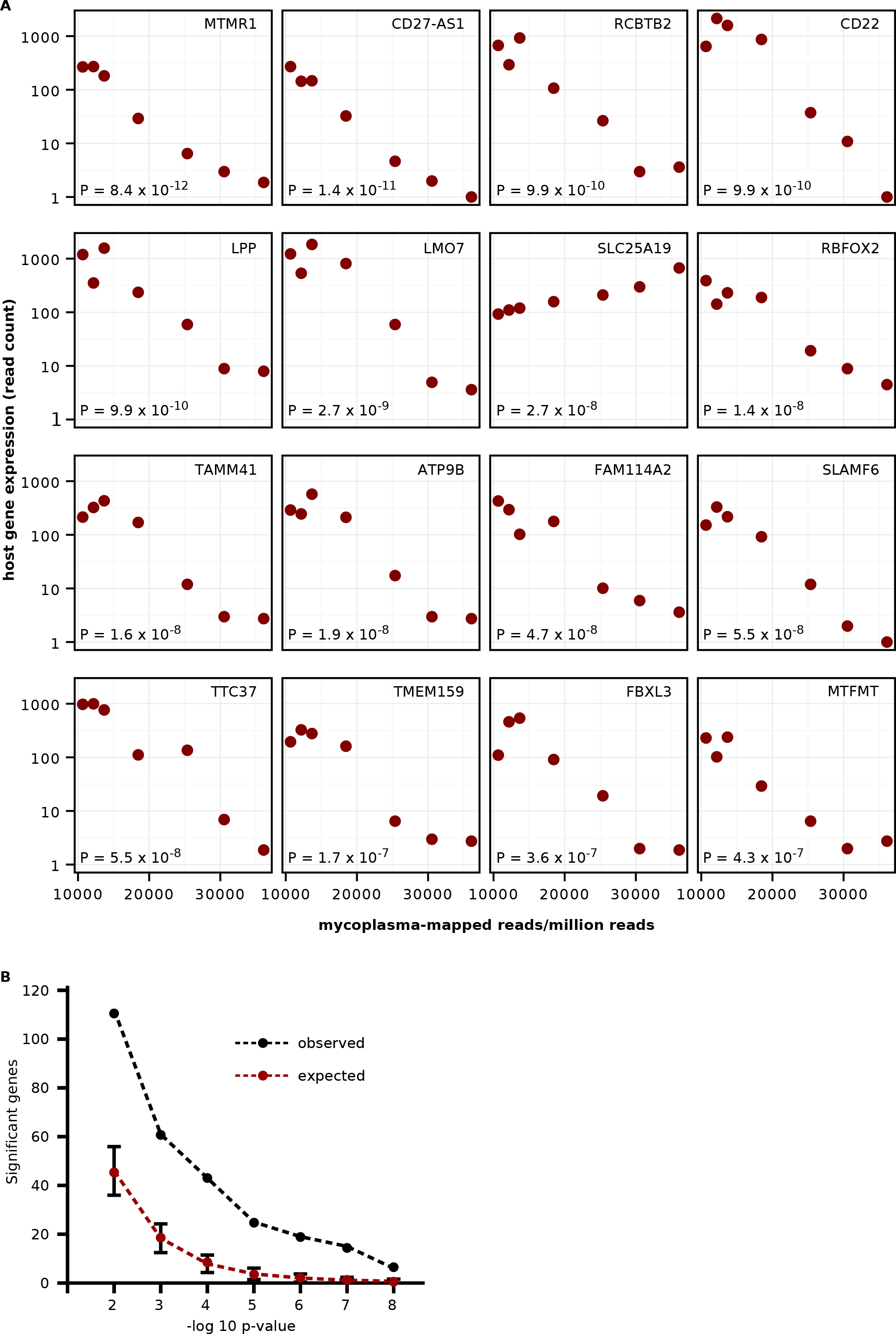
Association between host gene expression and mycoplasma-mapped reads from single-cell RNA-seq. A) Scatter plots of mycoplasma-mapped reads per million and host cell gene expression in single cell (DG-75) RNA-seq. Gene symbols are indicated in the upper right corner of the respective plots. P-values from the association test are indicated in the bottom left. B) To assess the likelihood of obtaining significant genes by chance, the mycoplasma counts were permuted 1000 times. The analysis was repeated with each permutation. The expected number of significant genes is plotted in red. The observed number of significant genes is in black. Error bars are standard deviations.

### Economic impact of mycoplasma contamination

To assess the potential financial impact of mycoplasma contamination, we first tallied the cost of NIH-funded research involving cultured cells. We queried the NIH Research Portfolio Online Reporting tools (RePORT) database for grants from 2013-2014 containing the keywords “cell culture” or “cell line”. A total of $3.17 billion in funded grants across the 27 NIH institutes and centers contained these terms (Figure 5). Using the contamination rate of 11% from our earlier results (Figure 2), we estimated that some portion of approximately $350 million of NIH-funded research is potentially affected.

**Figure 5.**
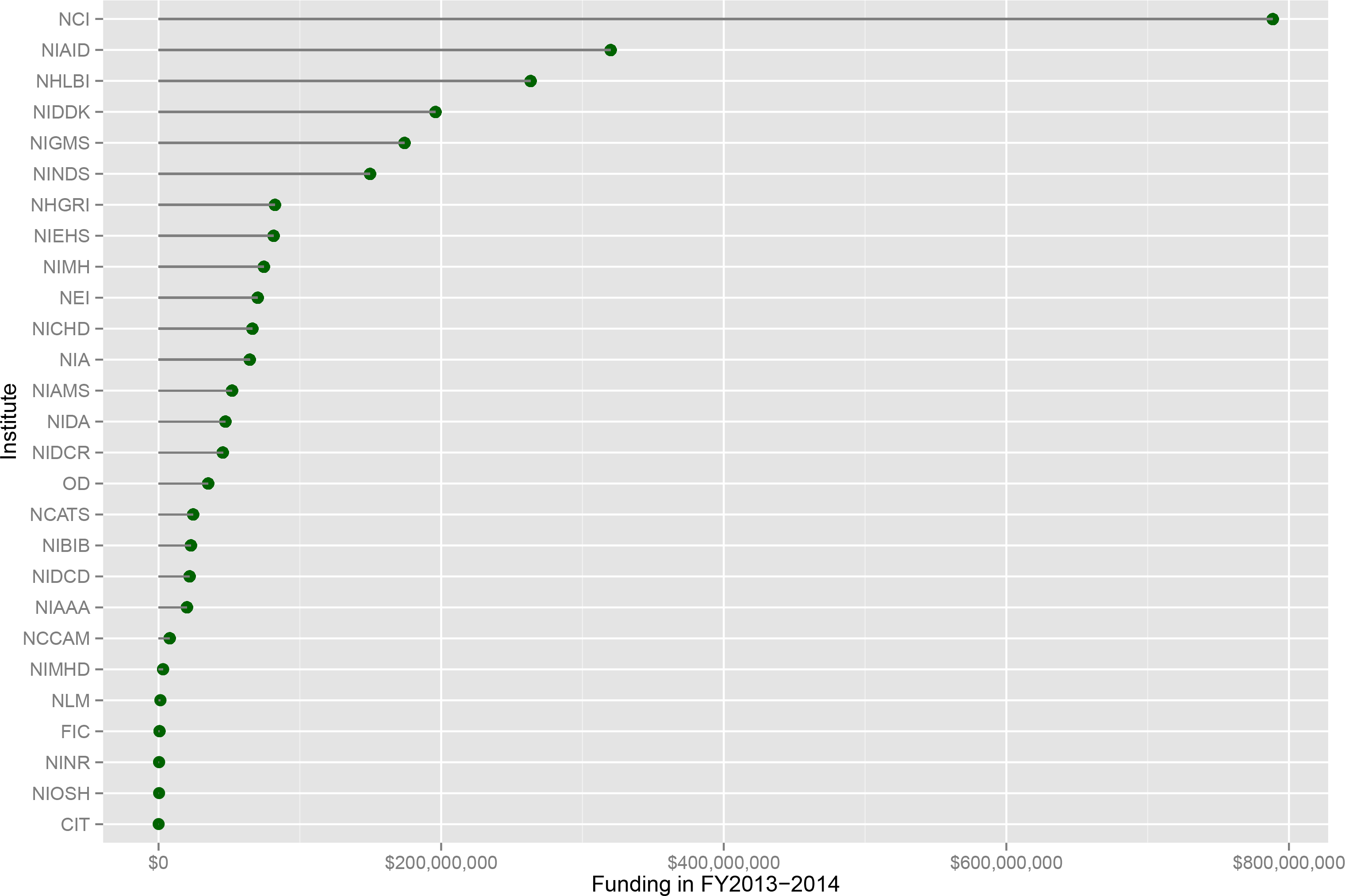
NIH-funded grants involving cell culture. The NIH RePorter was queried for grants from 2013-2014 that involved the culturing of cells. The results are broken up by the 27 NIH institutes and centers.

## DISCUSSION

This study was inspired by a recent incident of mycoplasma contamination in our lab. We wondered how often this occurred in other labs. There have been several such surveys over the last three decades (Rottem and Barile 1993; Coronato and Coto 1991; Drexler and Uphoff 2002). These studies used DNA fluorescent staining or PCR-based methods to detect mycoplasma contamination in collected samples. The contamination rates varied from 15% to 70%. Further, a recent analysis of DNA sequences from the 1000 Genomes project suggested 7% of the samples were contaminated (Langdon 2014). This study represents the first unbiased (and one of the most extensive) surveys for mycoplasma contamination. We leveraged publicly available RNA-Seq datasets in GEO to determine the prevalence of contamination today. Importantly, these entries were comprised of various sample types under different experimental conditions and originated from multiple institutions. We estimated that about 11% of GEO series were contaminated based on a cutoff of 100 mycoplasma-mapped RPM in one or more samples. However, this is likely an underestimate. For one, in our analysis, we discarded mycoplasma-mapped reads that were not unique to mycoplasma. Some of these included reads that mapped to other bacteria and hence could not be unambiguously associated with mycoplasma. While they may or may not be mycoplasma, they are bacterial. Secondly, Figure 1 suggests that a cutoff of 10 mycoplasma-mapped RPM was equally powerful to discriminate between cultured and non-cultured samples. At this cutoff, the contamination rate was closer to 15%. Nonetheless, no matter the exact rate, our study suggests mycoplasma contamination remains a significant and costly problem.

We also identified interesting characteristics of mycoplasma contamination in RNA-seq data. Notably, we found that the rate of mycoplasma contamination was similar for poly(A)- selected and non-poly(A)-selected series (Figure 2). This was unexpected because 1) 90% of the mycoplasma-mapped reads mapped to rRNA (Figure 1B) 2) Bacteria transcripts are degraded upon polyadenylation (Dreyfus and Régnier 2002) and 3) the mycoplasma genome lacks homologs for polyadenylating enzymes (Sarkar 1996; Portnoy and Schuster 2008). One explanation for this result is that the AT-rich mycoplasma genome (∼28% GC-content) may result in higher susceptibility of transcripts to oligo-dT mediated capture. This is perhaps a desirable outcome as contamination in future RNA-Seq experiments may be easily ascertained without masking from the enrichment method.

We mapped sequence reads to four complete mycoplasma genomes: *M. hominis*, *M. hyorhinis*, *M. fermentans*, and *A. laidlawii*. *M. hominis* and *M. fermentans* are associated with human diseases; the former, pelvic inflammatory disease, the latter, rheumatoid arthritis (Johnson et al. 2000; Taylor-Robinson 2007). *M. hyorhinis* is commensal in the respiratory tract of pigs (Kobisch and Friis 1996). *A. laidlawii* has a broad host range but is typically found in cattle. As such, *A. laidlawii* was linked to contamination of commercial bovine serum in the 1970s (Drexler and Uphoff 2002). These four species, along with *M. orale* and *M. arginini*, are found in 90-95% of mycoplasma contaminated cell lines (Drexler and Uphoff 2002). In our study, we found *A. laidlawii* was least represented in contaminated samples (Figures 2B-C, 3B-C). This may be indicative of the reduced likelihood of contamination from bovine serum as commercial products are now routinely 0.1 micron sterile-filtered.

Lastly, we identified 61 genes associated with mycoplasma contamination in a single cell RNA-seq dataset. Unfortunately, we could not identify series containing contaminated and non-contaminated samples. Therefore, we measured the association between the number of mycoplasma-mapped reads per sample and host gene expression in a contaminated series. Consequently, directionality of effect could not be determined. That is, differences in the extent of mycoplasma contamination may have driven host gene expression or differences in host gene expression (especially as it relates to variability of single cells) may have conferred resistance (relatively speaking) to contamination or impacted mycoplasma gene expression. It is also possible that the associations seen here are not biological per se, but instead indicative of some technical artifacts associated with the presence of the AT-rich mycoplasma transcripts in the sample. Nonetheless, this is equally problematic as it effects the interpretation of the results. Hence, further controlled studies are needed to fully understand the effect of contamination on the quality of RNA-Seq data and possibly how to account for this in data analysis.

To try to estimate the potential economic cost of this widespread contamination, we searched for grants that used the terms “cell culture” or “cell lines” in NIH RePorter. This analysis found > $3 billion dollars was invested in this research in 2013-2014. If we assume that the contamination rate for our set is the same as the average NIH sponsored lab, this puts some portion of ∼ $350 million at risk. We are heavy users of cell culture, with approximately 50% of our consumables budget going to these experiments. Other labs may be closer to 10%. So somewhere between dozens and ∼$100m are at risk. Put simply, the economic impact of this problem is significant.

Lastly, our study has broader implications for the analysis of high-throughput sequencing data. Too often in our analysis pipelines, we disregard unmapped reads as technical artifacts or attribute them to problematic or unsequenced regions of the reference genome. While this may be true most of the time, this study suggests such reads warrant further investigation—particularly when they constitute a large number of the total reads. We focused on mycoplasma in this study as a proof of concept due to its known prevalence. However, the methods used here are easily applicable to other known (and unknown) contaminants of cell culture including other bacteria, yeast and viruses.

## METHODS

All Perl scripts used in this study are available as supplemental files. Directions for their use are located in the README file. The BLAST analysis was performed through Amazon Web Services (8-core, 70GB of RAM). All other analyses were done on the high-performance computing cluster of the Penn Genomics Frontier Institute.

### Obtaining sequence files

GEO series IDs and descriptions for all RNA-seq experiments were obtained through the GEO DataSets Advanced Search Builder (http://www.ncbi.nlm.nih.gov/gds/advanced). Results were limited on the query parameter “DataSet Type” for "expression profiling by high throughput sequencing" and "non coding rna profiling by high throughput sequencing". The results were further filtered to include only primates (*Homo sapiens, Pan troglodytes, Macaca mulatta, Macaca fascicularis, Pan paniscus, Gorilla gorilla*) and rodents (*Mus musculus, Rattus novergicus*). SOFT files for all the GEO series were downloaded. From these files, the sample IDs were obtained and used to download the corresponding raw sequences using the fastq-dump utility of the SRA toolkit (http://eutils.ncbi.nih.gov/Traces/sra/?view=software).

### Identifying poly(A)-selected and cultured samples from sample descriptions

SOFT files were parsed and searched for keywords under certain headings. For example, poly(A) selection was assumed if the library source or extraction protocols contained the keywords “polyA” or “oligo-dT” and similar variants. A sample was considered cultured if its description contained the keywords “cell line”, “fibroblast”, “MEFs”, “ESC”, “immortalized”, “DMEM”, “RPMI”, “Ham’s”, “McCoy”, “cultured”, “passaged”, “propagated”, “grown” and similar variants of these keywords.

### Mapping sequence reads to mycoplasma genomes

The mycoplasma genomes used in this study were downloaded from NCBI genomes: *M. hominis* ATCC 23114 (NC_013511.1), *M. hyorhinis* MCLD (NC_017519.1), *M. fermentans* M64 (NC_014921.1) and *A. laidlawii* PG-8A (NC_010163.1). RNA-seq reads were mapped to these genomes with Bowtie using the default parameters (Langmead et al. 2009). If a read was paired-end, only the first read was used in the analysis.

### Filtering non-mycoplasma-specific sequences from bowtie results

All unique bowtie mapped reads were aligned to NCBI’s nucleotide (nt) database using standalone BLAST+ (http://www.ncbi.nlm.nih.gov/books/NBK1763/). The following parameters were used: -db nt -num_threads 8 -outfmt "7 std sscinames" -max_target_seqs 100. A bowtie-mapped read was considered unique to mycoplasma if: 1) the scientific name matched “Mycoplasma” or “Acholeplasma laidlawii” in the BLAST output and 2) the entry from 1) was the best BLAST hit (i.e. with the lowest E-value).

### Mycoplasma effect on host gene expression

The complete sequences for the 7 samples comprising series GSE49321were downloaded with the SRA toolkit. The sequences were aligned to the human genome (hg19) using STAR (Dobin et al. 2013) with the following optional parameters: --runMode alignReads --runThreadN 8. Read counts for each gene were determined using HTSeq with the following optional parameters: -s no-m intersection-nonempty. Gene expression (counts) was modeled as a function of mycoplasma-mapped reads using DESeq2.

## DATA ACCESS

The sequence files used in this study were obtained from the SRA at NCBI and are freely available for download from the source.

## ACKNOWLEDGEMENTS

We thank David Lipman, Don Preuss, and Eugene Yaschenko at NCBI for their advice on the best ways to query and download the large volume of sequencing data available in SRA. We also thank Nick Lahens for assistance working with AWS and his thorough review of the manuscript. This work is supported by the National Institute of Neurological Disorders and Stroke (1R01NS054794-06 to JBH), the Defense Advanced Research Projects Agency (DARPA-D12AP00025, to John Harer, Duke University), and by the Penn Genome Frontiers Institute under a HRFF grant with the Pennsylvania Department of Health.

## REFERENCES

1. Altschul SF, Gish W, Miller W, Myers EW, Lipman DJ. 1990. Basic local alignment search tool. J Mol Biol 215: 403–410.

2. Anders S, Pyl PT, Huber W. 2014. HTSeq A Python framework to work with high-throughput sequencing data. http://biorxiv.org/lookup/doi/10.1101/002824 (Accessed March 15, 2014).

3. Chanock RM, Hayflick L, Barile MF. 1962. Growth on artificial medium of an agent associated with atypical pneumonia and its identification as a PPLO. Proc Natl Acad Sci U S A 48: 41–49.

4. Coronato S, Coto CE. 1991. [Prevalence of Mycoplasma orale as a contaminant of cell cultures in Argentina]. Rev Argent Microbiol 23: 166–171.

5. Dobin A, Davis CA, Schlesinger F, Drenkow J, Zaleski C, Jha S, Batut P, Chaisson M, Gingeras TR. 2013. STAR: ultrafast universal RNA-seq aligner. Bioinformatics 29: 15–21.

6. Drexler HG, Uphoff CC. 2002. Mycoplasma contamination of cell cultures: Incidence, sources, effects, detection, elimination, prevention. Cytotechnology 39: 75–90.

7. Dreyfus M, Régnier P. 2002. The Poly(A) Tail of mRNAsBodyguard in Eukaryotes, Scavenger in Bacteria. Cell 111: 611–613.

8. Gill P, Pan J. 1970. Inhibition of cell division in L5178Y cells by arginine-degrading mycoplasmas: the role of arginine deiminase. Can J Microbiol 16: 415–419.

9. Hopfe M, Deenen R, Degrandi D, Köhrer K, Henrich B. 2013. Host Cell Responses to Persistent Mycoplasmas - Different Stages in Infection of HeLa Cells with Mycoplasma hominis ed. Y.A. Kwaik. PLoS ONE 8: e54219.

10. Huang DW, Sherman BT, Lempicki RA. 2009. Systematic and integrative analysis of large gene lists using DAVID bioinformatics resources. Nat Protoc 4: 44–57.

11. Johnson S, Sidebottom D, Bruckner F, Collins D. 2000. Identification of Mycoplasma fermentans in synovial fluid samples from arthritis patients with inflammatory disease. J Clin Microbiol 38: 90–93.

12. Kobisch M, Friis NF. 1996. Swine mycoplasmoses. Rev Sci Tech Int Off Epizoot 15: 1569–1605.

13. Langdon WB. 2014. Mycoplasma contamination in the 1000 Genomes Project. BioData Min 7: 3.

14. Langmead B, Trapnell C, Pop M, Salzberg SL. 2009. Ultrafast and memory-efficient alignment of short DNA sequences to the human genome. Genome Biol 10: R25.

15. Love MI, Huber W, Anders S. 2014. Moderated estimation of fold change and dispersion for RNA-Seq data with DESeq2. http://biorxiv.org/lookup/doi/10.1101/002832 (Accessed June 8, 2014).

16. Miller CJ, Kassem HS, Pepper SD, Hey Y, Ward TH, Margison GP. 2003. Mycoplasma infection significantly alters microarray gene expression profiles. BioTechniques 35: 812–814.

17. Paddenberg R, Weber A, Wulf S, Mannherz HG. 1998. Mycoplasma nucleases able to induce internucleosomal DNA degradation in cultured cells possess many characteristics of eukaryotic apoptotic nucleases. Cell Death Differ 5: 517–528.

18. Portnoy V, Schuster G. 2008. Mycoplasma gallisepticum as the first analyzed bacterium in which RNA is not polyadenylated. FEMS Microbiol Lett 283: 97–103.

19. Razin S. 2006. The Genus Mycoplasma and Related Genera (Class Mollicutes). In SpringerReference, Springer-Verlag, Berlin/Heidelberg http://www.springerreference.com/index/doi/10.1007/SpringerReference_3750 (Accessed August 7, 2013).

20. Razin S, Yogev D, Naot Y. 1998. Molecular biology and pathogenicity of mycoplasmas. Microbiol Mol Biol Rev MMBR 62: 1094–1156.

21. Robinson LB, Wichelhausen RH. 1956. Contamination of human cell cultures by pleuropneumonialike organisms. Science 124: 1147–1148.

22. Rottem S. 2003. Interaction of mycoplasmas with host cells. Physiol Rev 83: 417–432.

23. Rottem S, Barile MF. 1993. Beware of mycoplasmas. Trends Biotechnol 11: 143–151.

24. Rottem S, Naot Y. 1998. Subversion and exploitation of host cells by mycoplasmas. Trends Microbiol 6: 436–440.

25. Sarkar N. 1996. Polyadenylation of mRNA in bacteria. Microbiol Read Engl 142 (Pt 11): 3125–3133.

26. Taylor-Robinson D. 1996. Infections due to species of Mycoplasma and Ureaplasma: an update. Clin Infect Dis Off Publ Infect Dis Soc Am 23: 671–682; quiz 683–684.

27. Taylor-Robinson D. 2007. The role of mycoplasmas in pregnancy outcome. Best Pract Res Clin Obstet Gynaecol 21: 425–438.

28. Tully JG, Taylor-Robinson D, Cole RM, Rose DL. 1981. A newly discovered mycoplasma in the human urogenital tract. Lancet 1: 1288–1291.

29. Young L, Sung J, Stacey G, Masters JR. 2010. Detection of Mycoplasma in cell cultures. Nat Protoc 5: 929–934.

